# Functional Redundancy in Local Spatial Scale Microbial Communities Suggest Stochastic Processes at an Urban Wilderness Preserve in Austin, TX, USA

**DOI:** 10.1101/2020.06.25.171660

**Authors:** Justin Stewart, Amy Ontai, Kizil Yusoof, Teresa Bilinski

## Abstract

Empirical evidence supports selection of soil microbial communities by edaphic properties across large spatial scales, however; less is known as smaller spatial scales (e.g 10s-100s of meters). The goal of this research was to evaluate the relationship between ecosystem characteristics and bacterial community structure/function in soils across small spatial scales in an urban preserve. We employed 16s rRNA gene sequencing, community level physiological profiling (CLPP), and soil chemical analysis to address this goal. We found no significant relationship between gradients in soil characteristics and community structure/function. In contrast, *Acidobacteria, Bacteroidetes*, and *Nitrospirae* responded to variation in edaphic properties. Taxa exhibited a wide range in dispersal, supporting our finding of community wide differences in taxonomy. Furthermore, there was high metabolic diversity within the bacterial communities despite preferential metabolism of water-soluble polymers (Tween 40/80). Carbon substrate utilization patterns also suggest dominance of functional generalists. Pairwise comparison of carbon substrate utilization patterns indicates that there are high levels of microbial functional redundancy within soils across the sampling area. Lastly, we found that edaphic properties did not shape the overall community structure and/or function, and our analyses suggest that stochasticity may play a role in bacterial community assembly in soils with the local spatial scale of this research study.

**Graphical Abstract:** One Sentence Summary: Microorganisms at small spatial scales were functionally similar despite subtle differences in community composition.

## 1.0 INTRODUCTION

Soils harbor an incredible diversity of microorganisms and our current understanding of soil ecosystems is that the microbial biodiversity of soils rivals or exceeds that of diverse aboveground ecosystems(Wall et al., 2010). These communities drive food webs by contributing to the organic matter pool through their biomass (Schlesinger and Schlesinger, 1997), acting as a food source(Scheu, 2002), and by mediating critical soil processes such as organic matter decomposition and nutrient cycling (Lladó et al., 2017). As a result, soil microbial communities significantly affect above-ground communities through plant growth and plant diversity (van der Heijden et al., 2008; Schnitzer et al., 2011; Wagg et al., 2014).

Edaphic properties act as environmental filters that strongly affect the structure and function of soil microbial communities, as suggested by Bass-Becking tenet that “everything is everywhere, but the environment selects” (de Wit and Bouvier, 2006). Climate gradients appear to significantly affect the diversity and function of soil microbial communities, likely due to differences in air temperature, soil moisture, and plant community composition with increasing latitude (Garcia-Pichel et al., 2013; Staddon et al., 1998; Wu et al., 2009) and elevation (Bryant et al., 2008; Yang et al., 2014). Also, many studies have found significant differences in microbial community composition and/or diversity across environmental gradients at large spatial scales (Fierer and Jackson, 2006; Garcia-Pichel et al., 2013; Griffiths et al., 2011; Lauber et al., 2009; Xue et al., 2018).

Previous research has demonstrated that soil microbial community structure shifts across continental soil pH gradients (Docherty et al., 2015; Griffiths et al., 2011; Lauber et al., 2009; Xue et al., 2018), and that the relative abundances of specific bacterial taxa, such as *Acidobacteria*, respond strongly to differences in soil pH (Jones et al., 2009; Lauber et al., 2009). While pH may be an important control on microbial community structure and function at large biogeographic scales in which pH, this effect is likely attenuated at local scales. Instead, within a site or a region there are often stronger gradients in other environmental factors, such as vegetation type, carbon (Bååth et al., 1995; Bossio and Scow, 1998; Drenovsky et al., 2004), and nutrient availability (Fierer et al., 2012a; Frey et al., 2004; Ramirez et al., 2012).

Despite these accomplishments we still observe variability in the ways in which the structure and function of soil microbial communities respond to specific environmental parameters. Functional redundancy has been suggested to be responsible for this variability (Allison and Martiny, 2008a; Rousk et al., 2009). In addition, recent research has highlighted the potential for stochastic processes to influence the structure and function of microbial communities (Caruso et al., 2011; Stegen et al., 2012). For example, soil microbial community assembly has been shaped by neutral ecological processes following significant ecological disturbances, such as wildfires (Ferrenberg et al., 2013). Other ecological processes affecting community assembly, such as dispersal, drift and facilitation, are being acknowledged as important factors in the structuring of macrobial and microbial communities (Emerson and Gillespie, 2008; Kembel, 2009). It is likely that in many environments, community structure is influenced by both neutral and deterministic processes, although there is a need for further research on the ecological conditions which promote stochasticity (Caruso et al., 2011).

The goal of this research was to investigate whether gradients in edaphic properties over relatively small spatial scales (10-100’s of meters) strongly influence bacterial community structure and function. We hypothesized that at a small spatial scale, pH, moisture, and organic matter content would drive microbial community structure and function. A combination of bulk and chemical soil characterization, analysis of community level physiological profiles, and 16S rRNA gene sequencing was used to test these hypotheses. We did not find a statistically significant relationship between edaphic characteristics, overall bacterial community structure or function across the sampled locations. The majority of sites exhibited functional redundancy despite differences in taxonomic composition which are likely due to high dispersal. The abundances of key bacterial taxa responded to specific soil characteristics, suggesting that these soil parameters may have a role in the selection of these taxa. We also found evidence suggestive of stochastic processes in community assembly in that differences in soil characteristics did not correlate with differences in community structure and function. This research adds novel evidence for understanding of the local processes that govern the links between microbial community structure, function, and assembly within soil ecosystems.

## 2.0 MATERIALS & METHODS

### 2.1 Site Description

Wild Basin Wilderness Preserve is a 227-acre urban preserve located in Austin, TX (at 30.3103455°N, −97.8234956°W). Wild Basin is situated in a unique context in multiple respects. It is affected by the rapid urbanization occurring in and around Austin(Bureau, 2010). The preserve is bordered by Texas State Highway Loop 360 to the west, which is a corridor of active suburban development. Also, Wild Basin is located in the Edwards Plateau Ecological Region, which is characterized by large deposits of limestone, dolomite, shales, and sandstone, and the region is characterized by extremely alkaline soils. Although there has been relatively little alteration of plant communities by human activities over the past 50 years, in 1961 a high intensity wildfire burned approximately 60% of the woody vegetation across the preserve (Westlake Fire Department, Personal Communication). Thirteen of the 19 sampling sites in this study were located within the area that was affected by the fire, however since 1996, sites sampled in this study have hosted established woody plant communities.

### 2.2 Soil Sampling

Soils were collected between May 2016 through June 2017 at Wild Basin Wilderness Preserve at 20 long-term monitoring locations throughout the Wild Basin Wilderness Preserve (Supplemental Table 1). At each site three representative samples were collected within a 5m^2^ plot. For community level physiological profiling (CLPP) and microbial community composition, triplicate soil samples were homogenized, and a single composite sample was used for Ecoplate inoculation and DNA extraction. Methods for analyzing edaphic properties can be found in the supplemental.

### 2.3 Community Level Physiological Profiling (CLPP)

Microbial community function was measured using BiOLOG EcoPlates (BiOLOG, USA), 96 well plates consisting of different carbon guilds (carbohydrates, polymers, carboxylic acids, amines/amides, amino acids). These plates allow for the colorimetric measurement of metabolism through the reduction of a tetrazolium dye, which yields optical density (OD) data for each substrate. EcoPlates were inoculated with 100 uL of 10-^3^ diluted soil solutions and incubated in Multiskan FC Plate Reader (Fisher Scientific, USA) for 72 hours with absorbance at 595nm measured every half hour (Gomez et al., 2006).

### 2.4 Microbial Community Structure

Methods for 16s rRNA sequencing can be found in supplemental methods. Sequences have been deposited in the National Center for Biotechnology Information (NCBI) Sequence Read Archive (SRA) under SUB7125886.

### 2.5 Data Analysis

Statistics were calculated in R (3.6.0) using the *vegan* package (Oksanen et al., 2018) and graphics were produced using *ggplot2* (Wickham et al., 2008). Edaphic dissimilarity was measured using the Bray-Curtis metric. CLLP data were generated by averaging the triplicate absorbance values at 72 hours where the average absorbance value of the water wells was subtracted from the average of all other wells with any negative values being changed to zero. CLLP diversity was calculated using the Shannon Index (H) with the equation −Σ p(ln*pi), where pi is the proportion of the individual carbon substrate absorbance value relative to the total absorbance at the site(Shannon and Weaver, 1963).

Average Well Color Development (AWCD) was calculated as ΣOD_i_/n_(substrates)_ at 72 hours for each BiOLOG Ecoplate as a metric for total CLLP activity(Gomez et al., 2006). This equation was modified to ΣOD_(substrates in guild)_/n_(substrates in guild)_ for analysis of carbon guild metabolism. Pearson correlation coefficients were calculated between soil characteristics and CLLP values. Mann-Whitney tests with Bonferroni correction were used to identify pairwise significant differences in community function (Supplemental Table 2).

Edaphic properties, CLLP, and community were examined for normality using a Shapiro-Wilk test and quantile-quantile plots. Outliers were determined as being 1.5 times the interquartile range and then re-examined for normality using the method described above. Any site not displaying normality after removal of outliers had a log transformation applied to it, and if still not normally distributed, was removed from data analysis. Bray-Curtis dissimilarity distance matrices for edaphic properties and community composition at the OTU level were generated using the *vegan* package in R (Oksanen et al., 2018).

OTU counts were transformed into relative abundances for analysis. Mann-Whitney tests with Bonferroni correction were used to identify pairwise significant differences in community composition (Supplemental Table 3). OTU matrix using Bray-Curtis distance was made with relative abundances of OTUs at each site excluding 7.1 and 8.1 due to sequencing errors and then used to create a NMDS ordination. The role of bulk soil characteristics was evaluated using the a permutational correlation function in *vegan* where the variable is regressed against NMDS axis scores. Spatial autocorrelation was tested between OTU level and CLLP Bray-Curtis distance and Euclidean geographic distance matrices (Sokal and Wartenberg, 1983) using linear regression.

Community and functional difference graphs (Figures 7A,7B) were produced by taking the sum of pairwise Bray-Curtis dissimilarity values for each site and subsequent linear regression. Dispersal was measured by counting the presence of an OTU at a site and taking the sum of all sites. These values were then ranked and broken into 4 equal categories (high, moderate high, moderate-low, low).

## 3.0 RESULTS

### 3.1 Edaphic Properties

Soils were slightly alkaline, with average pH values of 7.64 ± 0.27. The range in soil pH was 7.18-8.26 with only two sites having a pH above 8.0 (sites 7.1, 9.1). Soil moisture was more variable, with an average value of 17.83% ± 6.0% and a range of 7.91%-30.71% across all sites. SOM had an average value of 12.51% ± 3.76%. None of the bulk soil characteristics were significantly correlated with soil temperature, depth, or elevation. Further characterization of edaphic properties can be found in Supplemental Table 4.

### 3.2 Community Level Physiological Profiling (CLPP)

Total metabolic activity in the soil samples, as measured through AWCD, ranged from 0.200-0.853. Overall CLPP profiles showed that polymers were the most used guild of carbon compounds followed by amines/amides, carbohydrates, amino acids, and lastly carboxylic and acetic acids (Figure 2). Absorbance values greater than 0.2 were measured for all of the carbon compounds in the Ecoplates for at every site/soil sample. A high degree of variability in substate use was found for every compound with the exception of D-Xylose and 2-Hydroxy Benzoic Acid which were used the least of all compounds (Figure 3). The most used compounds, Tween 80/40 (the most used substrate at 9/15 sites) demonstrated a wide range in use (the magnitude of absorbance values) (Figure 3B).

**Figure 1.**
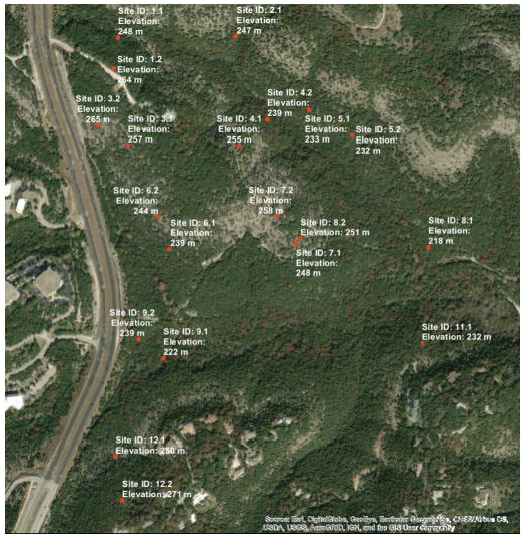
Map of Wilderness Preserve showing site locations and elevation, generated using a USGS DEM and ESRI ArcMap (10.4.1).

**Figure 2.**
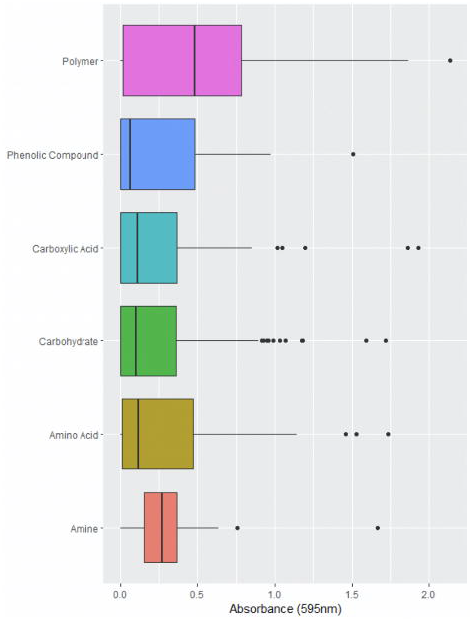
CLPP separated by carton guild from Biolog EcoPlates with average total absorbance (595nm) of all substrates in each guild at 72 hours of incubation.

**Figure 3.**
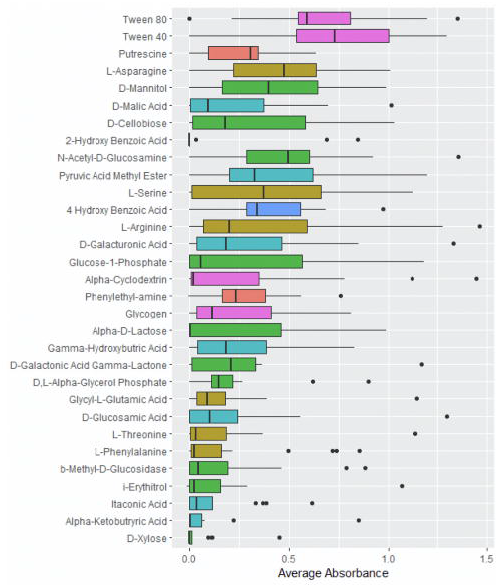
Functional diversity and abundance of substrate utilization a) across different sampling sites and colored by the most used carbon substrate; B) average total utilization (measured by absorbance at 595nm) of all substrates at all sites in after 72 hours of incubation.

A wide breadth of functional diversity was observed (Figure 3A) that varied spatially. Site 4.1 had the greatest diversity of carbon substrate use of all the sites while site 3.1 had the lowest carbon substrate use diversity. Comparison of all absorbances for across all compounds pairwise using Mann-Whitney tests across the sites revealed significantly different functional profiles of carbon substrate usage for 3 of the 19 sites (Supplemental Table 2). A mantel test between Bray-Curtis dissimilarity matrices of CLLP data and edaphic properties showed an insignificant relationship between CLLP profiles and soil characteristics (p > 0.05). Furthermore, seasonality was seen to not influence community function (PERMANOVA, R2=0.1573, p=0.555).

### 3.3 Microbial Communities

11 bacterial phyla were identified as the most abundant (> 5% on average) in all samples and phyla. *Proteobacteria* was the most dominant phylum and comprised over 33% of all reads with *Actinobacteria* also accounting for over 30% of sequence reads (Figure 4). All phyla were excluding those grouped into *Other* were present at all sites while their relative abundance differing. Phyla Shannon diversity ranged from 1.513 to 1.944 with an average 1.704 reported value. Spatial variation in community composition was found pairwise at 13 out of 16 sites (Supplemental Table 3). Microbes were largely either highly (31.08%, Figure 5) or lowly dispersed (45.77%). Seasonality did not not influence community structure (PERMANOVA, R2=0.19026, p=0.421).

**Figure 4.**
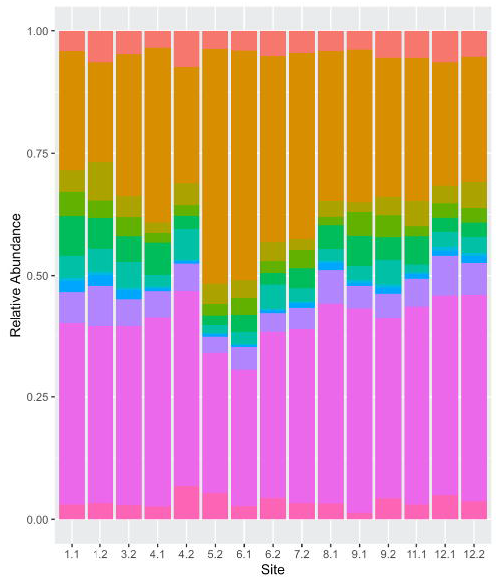
Relative abundances of phyla found at all sites excluding 2.1, 3.1, 5.1 and 7.1. Phyla with blow 0.5 % abundance were grouped into the “other” category.

**Figure 5.**
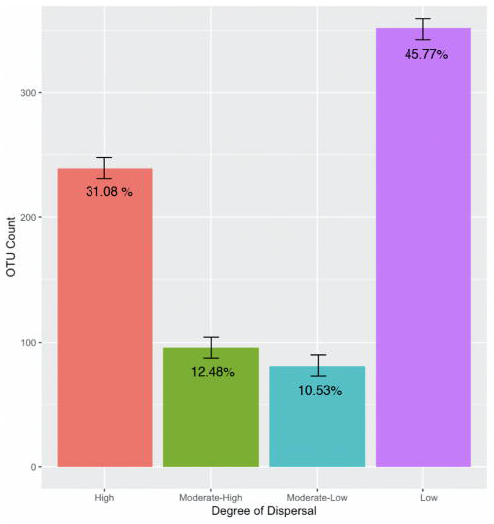
OTUs separated into dispersal categories with standard deviation bars and percentages of the community in the category. High: found at > 11 sites, Moderate-High: found at < 11 > 7 sites, Moderate-Low: found at < 8 > 4, Low: found at < 5 sites.

A mantel test demonstrated no significant relationship between the overall OTU level community composition and soil characteristics (r= 0.34, p > 0.05) or spatial autocorrelation (r= −0.12, p > 0.05). Despite these findings, specific taxa displayed deterministic patterns to edaphic properties.The relative abundances of *Acidobacteria* and *Nitrospirae* increased with soil moisture content (Pearson’s Rho= 0.63,p < 0.01), while the relative abundance of *Nitrospirae* decreased across the soil organic matter gradient (Pearson’s Rho = −0.69, p = 0.003). It was also observed that the relative abundances of *Bacteroidetes* increased with soil pH values (Pearson’s Rho = −0.49, p < 0.05) while the relative abundances of *Acidobacteria* decreased with increasing pH (Peason’s Rho= −0.52,p < 0.05). Based on a permutational correlation test (r = 0.031, p-value > 0.05, Figure 6B.) of dissimilarity in community composition (OTU level) and edaphic properties, edaphic variables did not structure the community as a whole. In addition, functional and OTU dissimilarity displayed the same pattern (Mantel Test, r=0.0231, p-value > 0.05)

**Figure 6.**
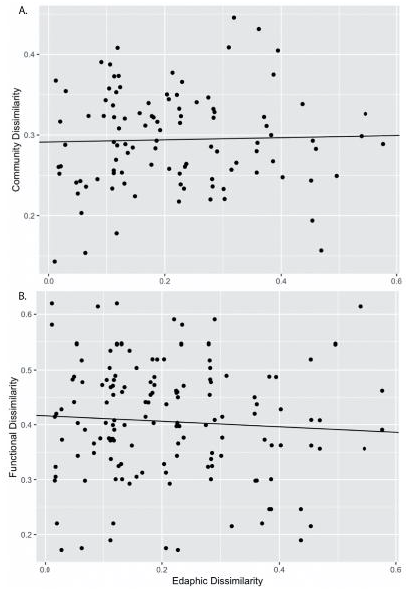
A: Difference in edaphic properties versus OTU level community difference. r = 0.031, p-value > 0.05, sites 3.1, 3.2, 5.1, 7.1, 7.2, 8.2, and 9.1 were removed from analysis due to lack of corresponding data between the sets. B: Difference in edaphic properties versus difference in function (CLPP profiles). y=0.05026x+0.41635, r=-0.066, p-value > 0.05. Sites 3.1, 3.2, 5.1, 8.2, and 9.2 were removed from analysis due to lack of corresponding data between sets.

## 4.0 DISCUSSION

The goal of this research was to test the relationship between bulk soil characteristics (pH, moisture, and organic matter content), and the structure and functioning of local microbial communities. Mantel tests showed there was not a statistically significant correlation overall between edaphic properties and either microbial community structure or function, despite that there was some variability in soil properties across sites, namely soil moisture and soil organic matter. We did not find evidence for significant differences between communities based on taxonomic structure however, it appears that the abundances of specific taxa (*Acidobacteria, Bacteroidetes, Nitrospirae*) are significantly correlated to specific edaphic properties. With regards to function, a wide range of carbon substrates were metabolised at all sites, but usage varied between the sites. The variation in function could not be explained by differences in soil properties. Furthermore, the communities were functionally redundant at the majority of sites despite significant differences in community composition at all but three sites. Together these findings provide preliminary evidence that local scale microbial community assembly may be subject to stochastic processes. This research has provided insight into the local importance of soil characteristics in structuring the taxonomic membership and functional capacity of these microbial communities. In summary, we have found that at the local scale these soil microbial communities experience a wide range in dispersal and exhibit functionally redundancy, despite subtle differences in taxonomic structure.

### 4.1 The relationship between environmental characteristics and microbial community structure

Multivariate microbial community structure as a whole was not determined by edaphic properties. Previous research has established soil pH as an important factor in the structure of microbial communities at the continental (Fierer and Jackson, 2006; Hartman et al., 2008; Lauber et al., 2009), regional (Griffiths et al., 2011; Hartman et al., 2008; Jenkins et al., 2009), and local scale (Baker et al., 2009; Philippot et al., 2009). However, we did not find evidence that soil pH significantly structured soil microbial communities, nor did we find a relationship between soil pH and bacterial community diversity in our study. This may be due to the buffering effects of carbonate and high calcium (Tavakkoli et al., 2015) in the underlying bedrock (limestone and dolomite) in the Edward’s Plateau (Barker et al., 1994). There may not be sufficient variation in pH for environmental filtering of the community at small spatial scales.

Aside from the lack of community wide response to soil pH, responses by individual taxa were observed. Our finding of *Bacteroidetes* abundance having a negative response to soil pH contrasts with the current literature (Lauber et al., 2009; Rousk et al., 2010). These studies take place at the continental scale (with stronger spatial gradients in the concentrations of electron donors and acceptors than in our study) and in laboratory manipulations and may not take into account responses of *Bacteroidetes* along attenuated gradients of pH.

Our findings do compliment well established literature showing negative correlations of *Acidobacteria* and positive correlations of *Nitrospirae* with pH (Fierer et al., 2007; Griffiths et al., 2011; Jones et al., 2009; Lauber et al., 2008). These results suggest that pH and soil moisture may have a more fundamental factor in determining the abundance of certain bacterial taxa, such as *Acidobacteria*, whereas for other taxa (e.g., *Bacteroidetes*) pH may be a proxy for other environmental co-variables that have a more profound impact on their ability to grow and compete in the soil environment. Furthermore, the relative abundances of Acidobacteria and Nitrospirae increased with soil moisture content, while the relative abundance of Nitrospirae decreased across the soil organic matter gradient. Soil moisture and soil carbon affected the structure and function of microbial communities in microcosms and across a climatic gradient (Brockett et al., 2012; Drenovsky et al., 2010). Aside from measured edaphic properties, the role of plant community composition (Myers et al., 2001; Reese et al., 2018; Sielaff et al., 2018; Yang et al., 2014) and soil texture (Bach et al., 2010) have been identified as factors in the structure of microbial communities.

### 4.2 CLPP profiles & edaphic properties

Our finding that community structure did not define how its members will function conflicts with other studies (Fierer et al., 2012b; Griffiths et al., 1998). This may be attributed to stochastic processes or lack of variation in edaphic properties to require functional niches. Although all carbon substrates were used by every soil microbial community in our study, there was a great deal of variation in terms of carbon substrate preference which indicates a wide range of metabolic diversity. Furthermore, many previous studies investigating the relationship between taxonomic and functional diversity took place at larger spatial scales where there were larger gradients in environmental characteristics. Previous research has found that enzymatic function in soils changes as spatial scale changes (ranging from soil horizon to landscape level) (Decker et al., 1999; Parsons et al., 1991; Šnajdr et al., 2008). It is possible that despite the wide range in both bulk soil characteristics and chemistry that the variation was insufficient to necessitate functional differences to increase fitness.

Discrete patterns in carbon substrate preference were observed as well as each substrate being utilized at every site, although to varying degrees of metabolism. In agreement with literature (Gryta et al., 2014; Reese and Maguire, 1969; San Miguel et al., 2007) we found that polymers Tween 40/80 were extensively used by the community; however, the amount to which these polymers were metabolized varied (Figure 3). These compounds are water-soluble surfactants consisting of partial fatty acid esters combined with ethoxylated sorbitan. Given that this structure is similar to ecologically common carbon compounds (fatty acids from organic matter) it is suggestive of communities dominated by microorganisms with generalist metabolisms. Furthermore this suggests the possibility of extracellular enzyme digestion (McShan et al., 2016; Skujiņš and Burns, 1976) before sequestration by microorganisms given its large size and the energetic preference for the diffusion of small molecules across cell membranes (Decad and Nikaido, 1976; Reese and Maguire, 1969).

In contrast, with significant differences in community structure, our results are suggestive of functional redundancy with regards to carbon substrate metabolism in that only 3 out of 19 sites had significantly different functional profiles. This disconnect between structure and function offers two possible interpretations. First, that taxa are equal in function due to low selection of the community, as supported by our finding that few taxons responded to variation in edaphic properties. Second, that the individual taxon may not be redundant but when combined with all the other microorganisms in the community result in the same function at the community level (Allison and Martiny, 2008b).

### 4.5 Potential for stochastic processes driving community assembly and function

At our study site, in which soils that are relatively homogeneous at local spatial scales, it is likely that environmental selection does not exert a strong influence on microbial community structure and function. By contrast, dispersal may be a dominant force in community assembly patterns, especially when the microbial taxa exhibit functionally redundancy. This may occur when a niche is left open that is then filled by the approximate 31% of the community that is highly dispersed. For example, in our research we found that 240 taxa were found at 75% of all sites. In contrast to community wide patterns, individual taxa that may be more sensitive to specific environmental conditions may be widely dispersed but their abundances would be more closely related to shifts in key edaphic properties. Our findings suggest that microbial community assembly may be driven by factors other than environmental selection, possibly stochastic processes like dispersal and drift, as proposed by (Nemergut et al., 2013) and shown in our Figures 7A,7B. Other studies have demonstrated this disconnect between microbial community structure and function and environmental conditions (Graham et al., 2016; Ramette and Tiedje, 2007; Zhou et al., 2008)). There is also evidence that certain environmental conditions, such as a high C:N in soils, and ecological disturbances like wildfire, are associated with greater stochasticity in microbial community assembly (Evans et al., 2017; Ferrenberg et al., 2013). In fact, a catastrophic wildfire in 1960 significantly altered the plant communities, and possibly also the soil properties, at our study sites.

Recent research has elucidated abiotic stochastic processes that influence microbial community assembly, such as historical contingency, dispersal, ecological drift, and selection (Allison and Martiny, 2008a; Dumbrell et al., 2010; Evans et al., 2017; Nemergut et al., 2013; Vellend, 2010) that may be occuring in this study. There is evidence that stochastic processes may dominate assembly in environments where there is not a strong link between community structure and community function. Furthermore, the assembly of communities in which functional generalists dominate, such as in our study, may not be as affected as strongly by environmental selection pressures (Langenheder and Székely, 2011). This may be due to a lack of variation in edaphic properties great enough to make functional differences relevant for fitness.

Discerning how microbial communities are structured and function is essential given their importance to biogeochemical cycling. In this study we explored the relationship between edaphic properties and the soil microbiome. A wide range in microbial function was observed and was redundant across the site; this contrasts with differences in taxonomy. Furthermore, our findings are suggestive of stochastic patterns of community assembly.

## FUNDING

This work was supported by the Dr. Allan W. Hook Endowed Wild Basin Creative Research Grant and the St. Edward’s University Biology Department.

## ACKNOWLEDGMENTS

We would like to acknowledge Cole Calderon, Emily Hooser, Ana Hernandez, and the Wild Basin Wilderness Preserve Staff for their assistance.

## Notes

### Competing Interest Statement

The authors have declared no competing interest.

