## Supplemental for "Functional Redundancy in Local Spatial Scale Microbial Communities Suggest Stochastic Processes at an Urban Wilderness Preserve in Austin, TX, USA"

Supplemental Methods:

*Soil Analysis:*

Triplicate soil samples were dried at 70ºC for 24 hours for gravimetric moisture analysis before shipment to the Texas A&M AgriLife Soil Facility in sterile glass tubes for measurements of soil chemistry. pH was determined using a 1:2 solution of dry soil to deionized water and allowed to settle for a minimum of 30 minutes and then the use of a hydrogen selective electrode. Soil organic matter (SOM) was determined using a combustion method. Concentrations of P, K, Ca, Mg, Na and S were measured using Mehlich III extractant and ICP-MS. Conductivity was determined using a 1:2 solution of dry soil to deionized water and allowed to settle for a minimum of 30 minutes and then the use of a conductivity probe. Nitrate-nitrogen (NO_3_-N) is extracted from soils using a 1 M KCl solution. Nitrate is determined by reduction of nitrate (NO_3_-N) to nitrite (NO_2_-N) using a cadmium column followed by spectrophotometric measurement. Elevation values were generated from points using a USGS digital elevation model (DEM) in ArcMap (Build 10.6.1, ESRI).

*DNA Sequencing:*

 	DNA from samples was isolated using the MOBIO PowerSoil DNA Isolation Kit. Concentrations of DNA were measured using a Nanodrop Spectrophotometer. Aliquots of DNA were then run on a gel to validate extraction. The 16S HV4 rRNA gene region was amplified in a 30-cycle PCR with the 515F (5’-GTGYCAGCMGCCGCGGTAA-3’) - 806R (GGACTACNVGGGTWTCTAAT) primer set and using StarTaq Plus Master Mix Kit (Qiagen, USA) under the following conditions: 94°C for 3 minutes, followed by 28 cycles of 94°C for 30 seconds, 53°C for 40 seconds and 72°C for 1 minute, and a final elongation step at 72°C for 5 minutes.

Sequencing was conducted at MR. DNA (Shallowater, TX, USA) on an Ion Torrent PGM following the manufacturer’s guidelines and subsequent analysis with an in house proprietary pipeline. To summarize, sequences were trimmed of primers and barcodes then reads < 150 bp were removed, sequences with ambiguous base calls and with homopolymer runs exceeding 6bp were also removed. Sequences were denoised, OTUs generated and chimeras removed. Operational taxonomic units (OTUs) were defined by clustering at 3% divergence (97% similarity). Final OTUs were taxonomically classified using BLASTn against a database derived from RDP II (http://rdp.cme.msu.edu) and NCBI (www.ncbi.nlm.nih.gov).

Supplemental Table 1. Sites sampled with soil temperature, soil core depth, elevation, and GPS coordinates.

| Site | Soil Temperature (ºC) | Core Depth (cm) | Elevation (m) | Latitude (N) | Longitude (W) |
| --- | --- | --- | --- | --- | --- |
| 1.1 | 21.5 | 8 | 248 | 30.31206 | -97.824811 |
| 1.2 | 20.1 | 8 | 264 | 30.311391 | -97.824941 |
| 2.1 | 21.8 | 8 | 247 | 30.312033 | -97.821844 |
| 3.1 | 25.3 | 6.8 | 257 | 30.3097 | -97.824626 |
| 3.2 | 23 | 10 | 265 | 30.31015 | -97.825389 |
| 4.1 | 22.8 | 6 | 255 | 30.30964 | -97.82181 |
| 4.2 | 23.4 | 8 | 239 | 30.310203 | -97.821085 |
| 5.1 | 19.4 | 8 | 233 | 30.310386 | -97.820025 |
| 5.2 | 19.4 | 8.5 | 232 | 30.309818 | -97.81894 |
| 6.1 | 25 | 7.5 | 239 | 30.307425 | -97.82364 |
| 6.2 | 25.2 | 7 | 244 | 30.308129 | -97.823898 |
| 7.1 | 25.6 | 9 | 248 | 30.307473 | -97.820424 |
| 7.2 | 25.7 | 9 | 258 | 30.308079 | -97.820887 |
| 8.1 | 23.9 | 7 | 218 | 30.30734 | -97.81705 |
| 8.2 | 23.9 | 7 | 251 | 30.30759 | -97.82031 |
| 9.1 | 23.9 | 8 | 222 | 30.305038 | -97.823846 |
| 9.2 | 24.3 | 8 | 239 | 30.305469 | -97.824468 |
| 11.1 | 24.7 | 9 | 232 | 30.305241 | -97.817271 |
| 12.1 | 19.3 | 8 | 250 | 30.302922 | -97.825115 |
| 12.2 | 19.2 | 7.5 | 271 | 30.301964 | -97.824969 |
